# AutoSmarTrace: Automated Chain Tracing and Flexibility Analysis of Biological Filaments

**DOI:** 10.1101/2021.01.19.427319

**Authors:** Mathew Schneider, Alaa Al-Shaer, Nancy R. Forde

## Abstract

Single-molecule imaging is widely used to determine statistical distributions of molecular properties. One such characteristic is the bending flexibility of biological filaments, which can be parameterized via the persistence length. Quantitative extraction of persistence length from images of individual filaments requires both the ability to trace the backbone of the chains in the images and sufficient chain statistics to accurately assess the persistence length. Chain tracing can be a tedious task, performed manually or using algorithms that require user input and/or supervision. Such interventions have the potential to introduce user-dependent bias into the chain selection and tracing. Here, we introduce a fully automated algorithm for chain tracing and determination of persistence lengths. Dubbed “AutoSmarTrace”, the algorithm is built off a neural network, trained via machine learning to identify filaments within images recorded using atomic force microscopy (AFM). We validate the performance of AutoSmarTrace on simulated images with widely varying levels of noise, demonstrating its ability to return persistence lengths in agreement with the ground truth. Persistence lengths returned from analysis of experimental images of collagen and DNA agree with previous values obtained from these images with different chain-tracing approaches. While trained on AFM-like images, the algorithm also shows promise to identify chains in other single-molecule imaging approaches, such as rotary shadowing electron microscopy and fluorescence imaging.

**Statement of Significance:** Machine learning presents powerful capabilities to the analysis of large data sets. Here, we apply this approach to the determination of bending flexibility – described through persistence length – from single-molecule images of biological filaments. We present AutoSmarTrace, a tool for automated tracing and analysis of chain flexibility. Built on a neural network trained via machine learning, we show that AutoSmarTrace can determine persistence lengths from AFM images of a variety of biological macromolecules including collagen and DNA. While trained on AFM-like images, the algorithm works well to identify filaments in other types of images. This technique can free researchers from tedious tracing of chains in images, removing user bias and standardizing determination of chain mechanical parameters from single-molecule conformational images.

## Introduction

Filamentous structures are ubiquitous in biology, from DNA and intracellular cytoskeletal components like actin, microtubules and intermediate filaments, to extracellular components like collagen. The mechanical flexibility of these structures can be intimately related to their performance, which can be altered by mutations or other chemical modifications that lead to changes in function correlating with disease (1-3). It is thus important to have standard means of assessing and quantifying changes in mechanics.

One of the most prevalent parameters used to characterize the mechanics of filamentous structures is the bending stiffness, which is generally described through the persistence length of a filament. The persistence length can be obtained by imaging individual filaments and determining the length scale over which thermal fluctuations tend to randomize the direction of the filament’s backbone. For semiflexible filaments such as DNA and molecular collagen, the persistence length is on the order of 10-100 nm (4-7), and thus atomic force microscopy (AFM) has become the technique of choice for conformational imaging, as it has sufficient spatial resolution to resolve the contour of the backbone of these filaments (1-2 nm in diameter). For thicker, more rigid filaments, fluorescence microscopy can be used to image fluctuations of filament backbones (8, 9). Once spatial coordinates of the backbone are determined, statistical treatment of the directional correlation and of the through-space separation of points along the backbone can be used to determine mechanical parameters such as persistence length and inherent curvature (4, 6, 8, 9). Although changes in persistence length can indicate molecular alterations, at present image analysis presents a bottleneck to assessing molecular flexibility in a high-throughput manner.

At present, there is no standard technique for determining backbone coordinates from AFM images. Manual chain tracing is frequently done when image sets are not too extensive (5), but for larger numbers of images, manual tracing can become cumbersome. Furthermore, it is possible that there is variation in backbone coordinates selected by different users, which could lead to biased results. For these reasons, a number of different approaches for automating chain tracing have been developed (6, 10-13). Most of these involve an initial user input step to manually identify points along the chains (6, 11, 12); following the initial identification of points on or near the central line of the backbone, various approaches are implemented to automatically refine the backbone contour. Very few algorithms exist that provide fully automated chain detection and tracing (10, 13). We have found one of these algorithms (10) to be sensitive to background noise amplitude and correlation, and problematic for chains exhibiting irregular intensity along the contour (14). Thus, user intervention may be required even in an automated routine to exclude erroneously detected traces.

To date, machine learning has proven to be a powerful tool for microscopy in biological fields, and it can improve significantly the efficiency and performance of large-scale image analysis, such as with medical diagnostics (15) and cellular imaging (16). More recently, deep learning techniques have been applied to imaging at the single-molecule level (17) to extract molecular parameters (18), improve localization (19), and automate analysis (20). A neural network thus presents a good candidate for the automation of the chain tracing process.

Here, we implemented a machine learning approach to automate the detection of chains in AFM images. Following pixel classification by a neural net, image processing is performed to determine an initial guess of the backbone of each chain. These coordinates are passed to our previously developed chain-tracing software SmarTrace (6, 14), which refines the contour and outputs coordinates used for determination of the persistence length. We validate this software, dubbed AutoSmarTrace, on simulated chains. We demonstrate its performance on experimental images by extracting persistence lengths from biological samples (DNA and collagen) that possess different persistence lengths, in some cases display inherent curvature or have a large globular end-domain, and whose images – recorded in different laboratories – contain various levels of background noise. We furthermore show the ability of the machine learning portion of the algorithm to identify filaments in electron microscopy and fluorescence images, which exhibit different appearances and noise profiles than AFM images. In all cases, AutoSmarTrace matches the performance of manually curated image tracing, offering a much more efficient route to chain identification and tracing for future applications.

## Methods

AutoSmarTrace is written in MATLAB (21). The program, documentation and example images are available at https://github.com/FordeLab/AutoSmarTrace. An overview of the image processing is provided in Figure 1. Simulated AFM images of filaments were created to train the neural network, and validation of the algorithm was tested on additional simulated images of chains of known persistence lengths that formed the “ground truth” for comparison.

**Figure 1.**
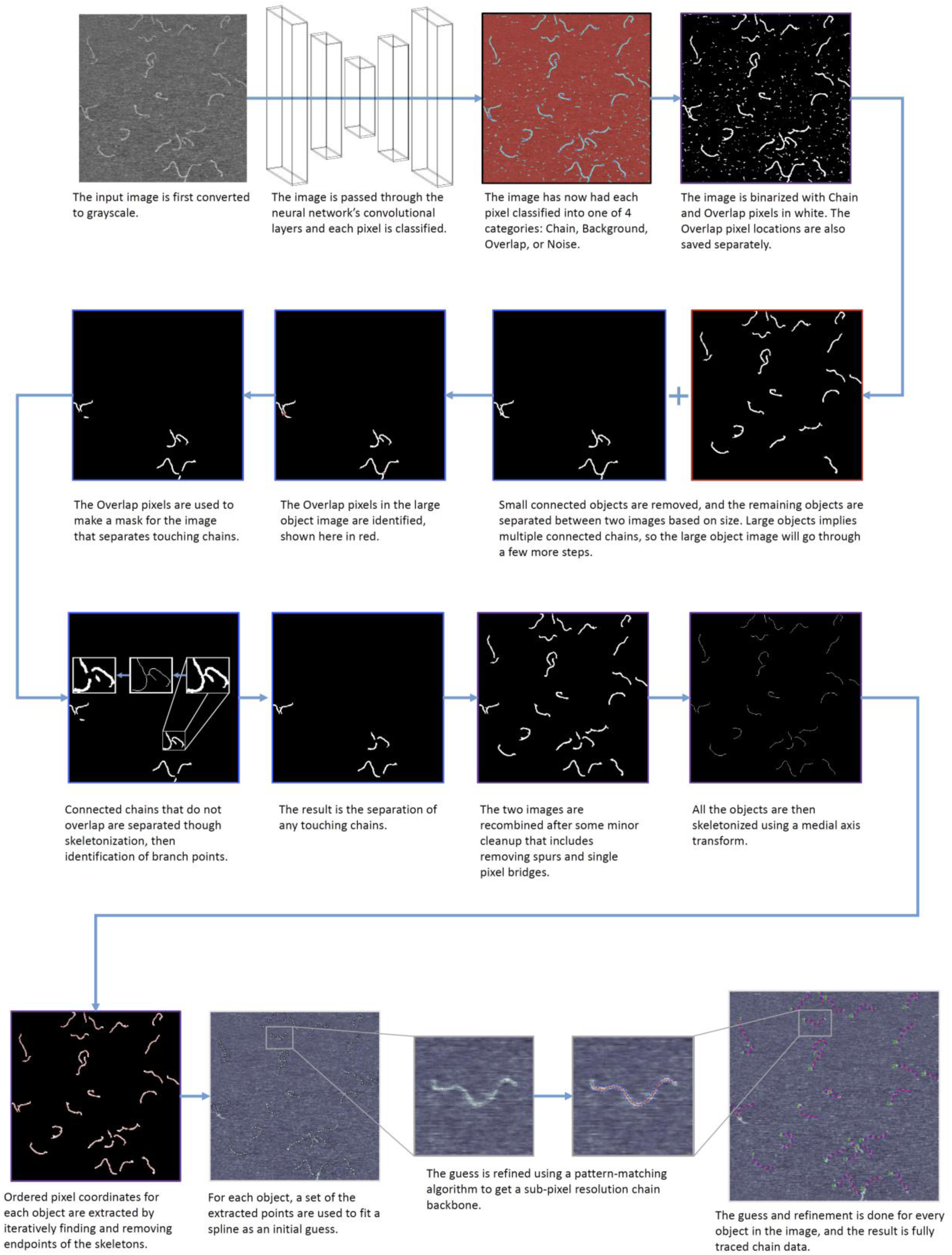
Workflow for AutoSmarTrace backbone identification and contour tracing algorithm.The final step (bottom line) of contour tracing is performed by the SmarTrace algorithm (6).

### Training set of images

The training set consisted of over 8000 simulated images of chains (contour length 300 nm), which contain varying numbers of chains, chain persistence lengths, chain widths and levels of background noise. Each image is 512×512 pixels in size with a corresponding pixel size of 4 nm, typical of AFM settings for imaging (6). The persistence length of chains varies between images from 20 nm to 200 nm at 5 nm intervals with an equal number of images per persistence length. This range encompasses experimentally measured persistence lengths for a variety of semiflexible polymers, including DNA, collagen and tropomyosin (4-7, 22). Chain width varied between images, with a value between three to six pixels. Background noise varies from zero to comparable to signal amplitude (such that the chains are not visually apparent in the images).

Each simulated image was generated by a previously described worm-like-chain-simulating algorithm (6). Chain parameters specifying persistence length and contour length were used to generate points along a two-dimensional chain backbone; 2D Gaussian distributions of a given width, centered on each point on the backbone, were summed to produce the final chain contour. AFM-like background was then added to the image by correlating Gaussian random noise through the Cholesky decomposition of a reference AFM image of background.(23) In some images, bright elliptical shapes were added to mimic larger pieces of dirt or debris that can appear in AFM imaging.

### Neural Network

The first component of the tracing algorithm is a convolutional neural network for the purpose of individual pixel classification (semantic segmentation) developed in the deep learning package of MATLAB Release 2018b (21). The layer architecture follows a downsample-upsample design with two sets of convolutional and transposed convolutional blocks of layers. Each downsampling block consists of a convolution, followed by a rectified linear unit (ReLU) layer, and then a max pooling. The final layers are for classification of pixels using a softmax layer into a cross-entropy loss layer. This final classification layer classifies each pixel as belonging to one of four categories: background, chain, overlap, and noise. The network was trained on the simulated images until an accuracy of 98-99% correct pixel identification was maintained.

### Image Processing

After the image has been segmented using the neural network, it is processed to provide backbone coordinates to a chain-tracing algorithm. This image processing part of the algorithm is also implemented in MATLAB (21). First, the classified image is converted to a binary image by equating the background and noise categories to black pixels and the chain and overlap categories to white while keeping a copy of the original labelled image. The copy is used to make another binary image containing only the overlap pixels in white. Both binary images have an area thresholding applied to each 8-connected object to remove small, misidentified groups of pixels, and then the overlap image is used to create a mask for overlapping regions of chains. The binary chain map is then cleaned further by removing highly circular objects, spur pixels, and single pixel bridges.

Further separation of connected chains not associated with overlap is done through the process of skeletonization. This procedure reduces objects down to a single-pixel-width, minimally connected skeleton by calculating a distance transform. The skeleton is then the pixels that correspond to the creases of zero curvature of the transform. This is done using the bwskel MATLAB function which utilizes the medial axis transform. A copy of each object is skeletonized, then any objects with branch points, pixels of the skeleton that have more than 2 connected pixels, are identified, and an area around the branch points are masked out in the original image. After each object in the image has undergone this procedure, there should be complete separation of connected chains.

To generate an end-to-end ordered list of pixel coordinates for each object, the skeletonization procedure is used once more. Each object is transformed, and the endpoints of each skeleton are identified. Their coordinates are saved (with one pixel arbitrarily chosen as the beginning of the chain) before they get removed from the skeleton. The process is then repeated with the new shorter skeleton until the ordered list is complete. Once each chain has had its points extracted, the lists are transferred to the SmarTrace algorithm to provide an initial spline that will be refined to a smooth backbone.

### SmarTrace

The SmarTrace program is chain-tracing software written in MATLAB that uses a pattern-matching algorithm to obtain a subpixel resolution spline for a chain backbone (6, 14). Prior to the development of the automated algorithm, SmarTrace used a graphical interface to allow users to zoom onto a chain, define points along the backbone, trace the chain, save the data, and move on to the next chain. In AutoSmarTrace, this is replaced by an automated system that reads all images in an input directory to a temporary datastore, then sequentially traces each image and saves the chain data as well as the traced image. The lists of points output from the image processing step replace the user-defined input points in the tracing process.

These initial points are then fit to a cubic spline with points 1 nm apart to serve as an initial guess. The guess is then refined by the pattern-matching algorithm to get a best fit for the chain backbone, resulting in a smooth sub-pixel resolution spline.

### Persistence length and curvature determination

Persistence length is determined as previously described (6). First, the splines across all images for a set of chains are pooled and randomly segmented into pieces of contour length 10-200 nm in 10 nm intervals. The segmentation is performed so that none of the resulting segments overlap with each other. Chains are resampled, and for each segment of each chain, cos*θ*(*s*) and *R*^2^(*s*) values are computed from the tangent vectors at the end points (Fig. 2). From all segments of a given contour length *s*, the mean and standard error of cos*θ*(*s*) and *R*^2^(*s*) are determined. These are fit with the following relations from the 2D worm-like chain model:

**Figure 2.**
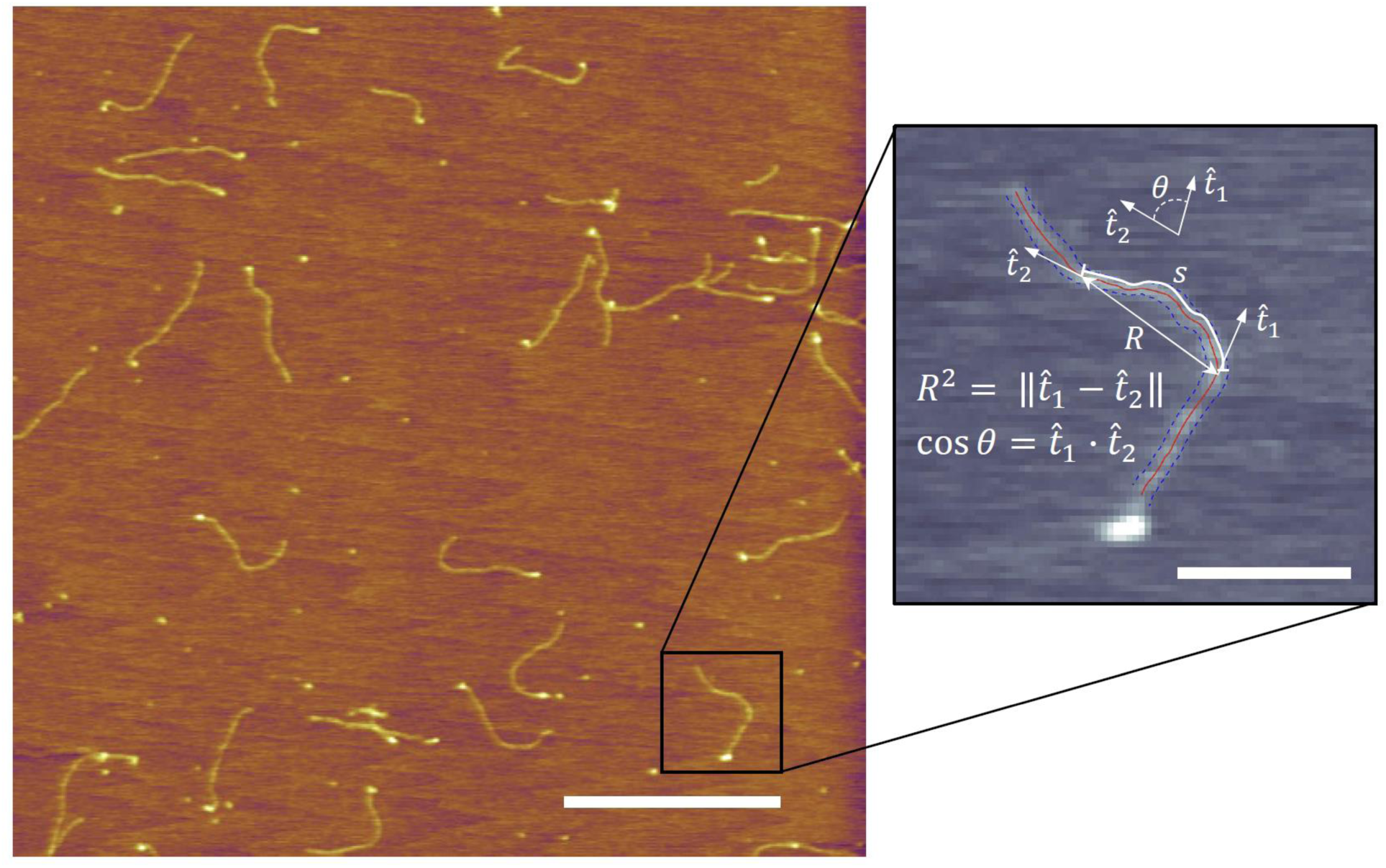
AFM image of pN-III collagen molecules. The two metrics used in our analysis are shown for two points separated by a contour distance of *s*: end-to-end distance squared, *R*^2^, and correlation of unit tangent vectors, 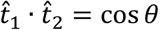. Scale bar in main image is 500 nm. Scale bar in zoomed image is 100 nm

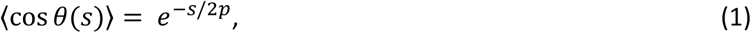

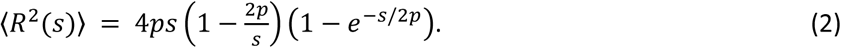

From these fits, the persistence length *p* is determined.

In some cases (e.g. collagen deposited under low ionic strength conditions; A-tracts of DNA; tropomyosin), chain conformations exhibit intrinsic curvature (6, 22, 24). Such conformations can be described by the curved WLC model, where the above relations are modified by the intrinsic curvature κ (6):

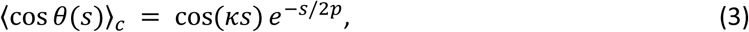

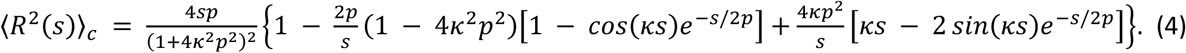

### AFM imaging

Bovine pro-N collagen III (pN-III) was a gift of Yoshihiro Ishikawa of the University of California, San Francisco, and was extracted from fetal bovine skin following a previously published protocol (25). Collagen was diluted into a solution of 1 mM HCl, 100 mM KCl at a final concentration of 0.2 μg/mL, where 50 μL of the diluted sample was deposited and allowed to incubate for 20 seconds on freshly cleaved mica (Highest Grade V1 AFM Mica Discs, 10 mm, Ted Pella). The excess unbound proteins were removed by rinsing with ultra-pure water, and the mica was then dried using filtered air. All proteins were imaged under dry conditions, and the solution conditions of the samples refer to the conditions in which they were deposited onto mica. Imaging was done with an Asylum Research MFP-3D atomic force microscope using AC tapping mode in air. AFM tips with a 160 kHz resonance frequency and 5 N/m force constant (MikroMasch, HQ: NSC14/AL BS) were used. AFM images of the other collagen samples were similarly obtained: rat type I collagen images and collagen types I and III deposited from 20 mM acetic acid are from reference (6), while type IV collagen images are from reference (7).

AFM images of DNA recorded in solution conditions were generously provided by Patrick Heenan and Tom Perkins of JILA and the University of Colorado Boulder (5).

### Electron microscopy imaging

Electron microscopy images of pN-III collagen were recorded and generously provided by Douglas R. Keene of the Micro-Imaging Center, Shriners Hospitals for Children, Portland, Oregon (26). Briefly, 5 mg of pN-III was resuspended in 1 ml of 200 mM acetic acid, which was diluted 1:5 in 1 mM HCl with 100 mM KCl. 30 μL of this 100 μg/ml collagen sample was mixed with 70 μL of glycerol and nebulized, using an airbrush, onto freshly cleaved mica. The sample was dried in vacuum and rotary shadowed using a Pt-C electron beam gun angled at 6 degrees relative to the mica surface within a Balzers BAE 250 evaporator. The replica was backed with carbon, floated onto distilled water and picked up onto 600 mesh grids. Photomicrographs were taken using an FEI G2 TEM operated at 120KV using an AMT 2Kx2K camera.

## Results

We have developed an automated process, dubbed AutoSmarTrace to trace chains in image data using a convolutional neural network to classify pixels, image processing to skeletonize chains, and supplied these coordinates to the previously developed SmarTrace algorithm to trace them with sub-pixel resolution. Figure 1 provides a summary of the steps involved in this automated process, which are described in more detail in the Methods section.

Key to this process is the classification of pixels by a trained neural network. The network in AutoSmarTrace was trained using 8000 simulated AFM images of chains with persistence lengths ranging from 20-200 nm at a variety of surface coverage densities, particulate “noise” contaminants, and background noise with amplitudes ranging from zero to comparable to signal levels from the chains. The network was trained to classify each pixel as belonging to a chain, to the overlap of chains, to noise or to background. Pixels belonging to chains were subsequently subjected to filtering algorithms to refine the choice of pixels belonging to extended regions of chains appropriate for analysis of chain flexibility. A skeletonization process was used to determine an ordered list of chain coordinates that were passed to SmarTrace as the input guess for the backbone. SmarTrace uses pattern-matching to refine the input coordinates and produce contour coordinates of each chain with sub-pixel resolution (6).

Visually, the tracing of contour backbones by AutoSmarTrace looks successful (e.g. final panel of Fig. 1). To test the performance quantitatively, we analysed the traced chains, determining their flexibility and, where appropriate, intrinsic curvature. The algorithm was tested on both simulated and experimental data to check for consistency and accuracy.

### Validation on Simulated Chains

First, we validated the algorithm on simulated chains with prescribed flexibility, which provided the ground truth. Large sets of images of chains with defined persistence lengths were simulated, then were traced. The set of images used for tracing here are a separate set from the training images, and around 200 μm of total contour length of simulated chains was traced for each persistence length determination. Simulated chains had persistence lengths ranging from 20 nm to 200 nm at 5 nm intervals. Tangent correlations ⟨cos*θ*⟩ and mean-squared distances ⟨*R*^2^⟩ were determined for different segment lengths from the traced contours; the dependence of these quantities on segment length were used to find the persistence length by fitting to equations (1) and (2), respectively. Persistence lengths determined from this tracing and analysis were compared to the ground truth persistence lengths that were defined by the simulation parameters.

As seen in Figure 3, AutoSmarTrace is able to accurately determine persistence lengths of simulated chains with persistence lengths from approximately 30-200 nm. For more flexible chains, the algorithm overestimates the persistence length. We provide reasons for this in the Discussion. It is also apparent from Fig. 3 that for persistence lengths below ∼110 nm, analysis of mean squared end-to-end distance (eq. 1) tends to overestimate the persistence length compared with analysis of tangent correlations (eq. 2). This was also found for manual usage of SmarTrace. Indeed, the mean squared end-to-end distance provides a good estimate for flexibility only for more stiff chains. In the tests on experimental data sets that follow, we report only the tangent correlation estimate for this reason.

**Figure 3.**
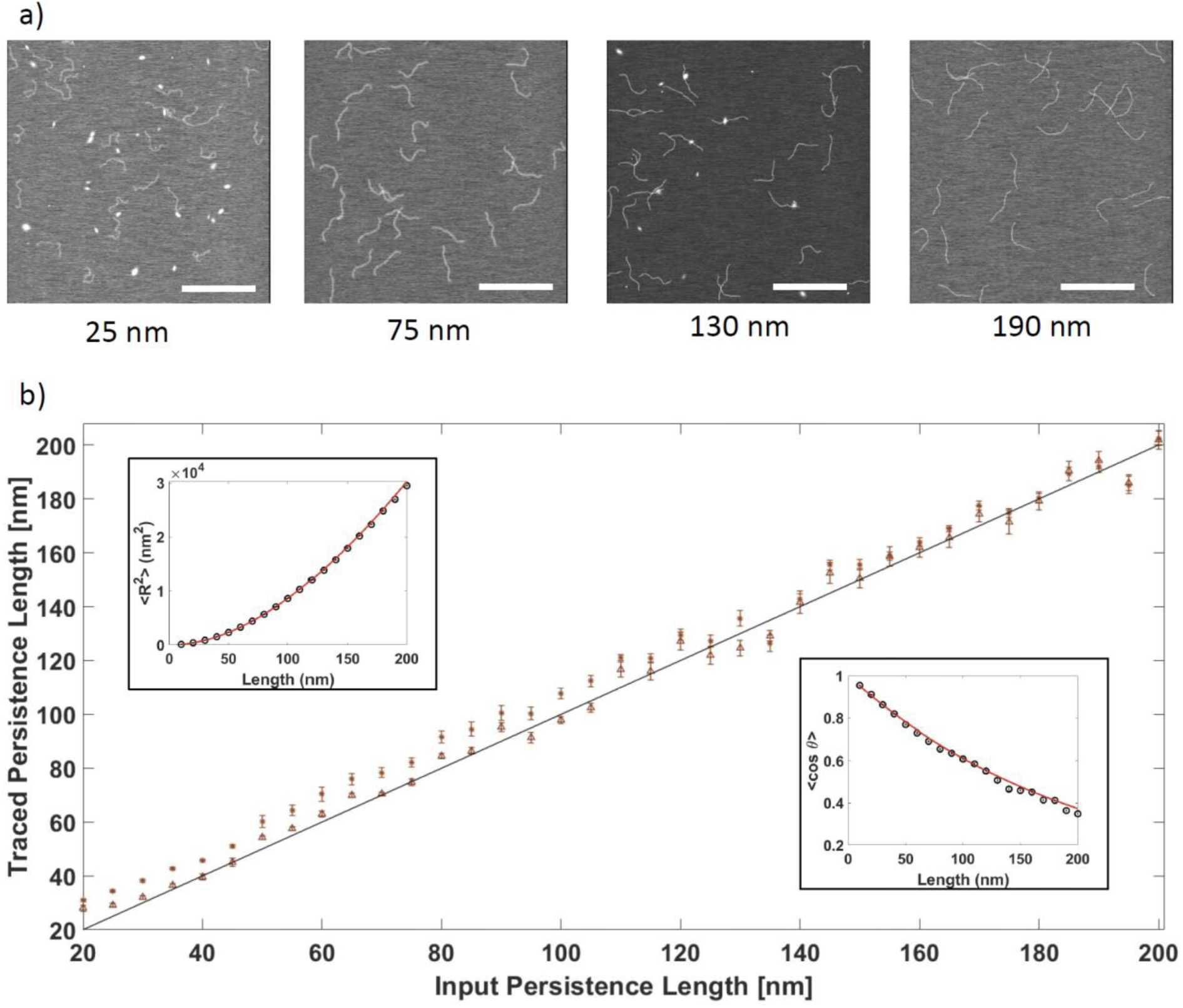
Performance of AutoSmarTrace on simulated AFM images of chains. a) Examples of simulated images of chains with different persistence lengths. Noise level, chain thickness, and the number of chains also varies between images. b) Values of persistence length obtained from analysis on splines from AutoSmarTrace compared to the input for the simulated chains. Shown are results for both ⟨cos*θ*⟩ (triangles) and ⟨*R*^2^⟩ (dots) fits, using equations (1) and (2), respectively. Insets show example fits for simulated chins with a persistence length of 50 nm. Error bars in the main panel are 90% confidence intervals from the WLC fits, and each point is determined from approximately 200 μm of traced contour length. Scale bars in images are 500 nm.

### Performance on experimental AFM images

To assess the performance of AutoSmarTrace on experimental images, we used it to trace chains in multiple sets of AFM images of different collagen samples. The resulting traces were analyzed to determine persistence length, and these results were compared to the values obtained through manual usage of the SmarTrace program. Additionally, we also traced a sample of AFM images of DNA with the automated and manual methods. These results were both compared to the value of persistence length independently obtained through a separate method of manual tracing (5).

Comparison of AutoSmarTrace with manual usage of SmarTrace and another manual tracing procedure provide comparable estimates of persistence length of these different samples (Figure 4a). These findings demonstrate the ability of AutoSmarTrace to determine chain flexibility in an unsupervised fashion, using AFM images recorded in either solution or dry conditions.

**Figure 4.**
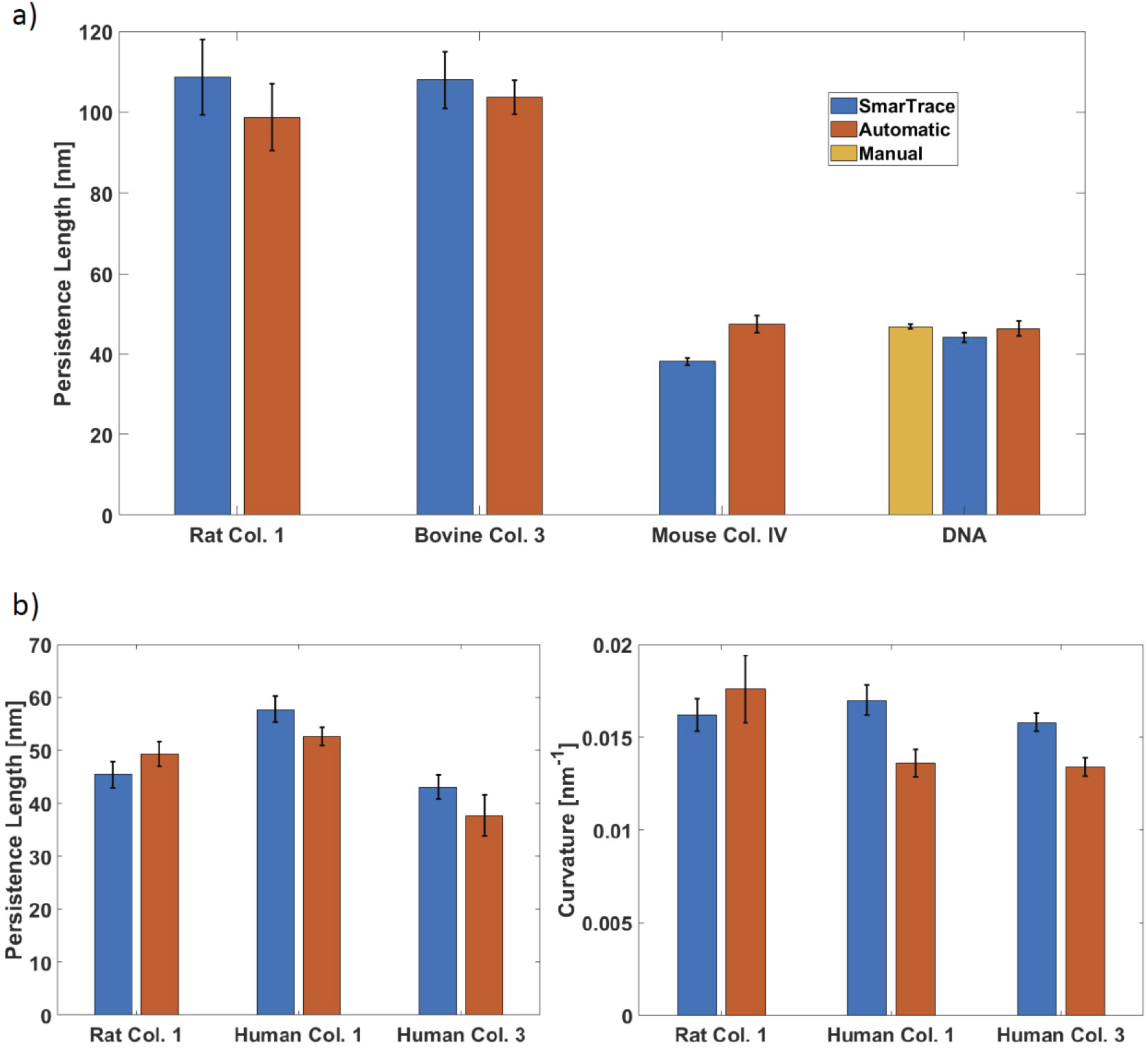
Performance of AutoSmarTrace on experimentally obtained AFM images. a) Persistence lengths obtained through analysis on spline sourced from manual usage of SmarTrace compared to running of AutoSmarTrace for several different polymer samples. Error bars represent 90% confidence intervals from the WLC fit. The DNA sample has an additional result from a manual spline fitting to points placed by hand with no further refining, as presented in (5). b) Persistence lengths and curvatures obtained through analysis on spline sourced from manual usage of SmarTrace compared to running of AutoSmarTrace for curved WLC samples. These curved WLC samples of collagen were deposited from 20 mM acetic acid (6). Error bars are 90% confidence intervals from the curved WLC fit.

Some biological filaments have been found to exhibit intrinsic curvature (6, 22, 24). We tested AutoSmarTrace’s ability to trace collagen chains deposited in conditions that exhibit curvature (6). Figure 4b shows the results of this analysis, obtained by fitting the measured tangent correlations ⟨cos*θ*⟩ and mean-squared distances ⟨*R*^2^⟩ with equations (3) and (4), respectively. Estimates of persistence length and curvature agree reasonably well between AutoSmarTrace and the manual usage of SmarTrace. In cases where the persistence length is underestimated by AutoSmarTrace (compared with manual SmarTrace), the curvature is also underestimated, likely due to a compensating effect arising from the inverse contributions of these two parameters to chain compactness in a curved worm-like chain.

### Performance on other experimental images

AutoSmarTrace complements a recently developed machine-learning algorithm for chain characterization, TopoStats, which was designed to determine contour length and topology of DNA (13). While both approaches were developed for analysis of AFM images, we were curious to see whether AutoSmarTrace could identify chains in images acquired with other approaches. We tested its performance on electron microscopy (EM) images of collagen and on fluorescence images of actin filaments in solution. Given the significantly different appearance of filaments and of background in these images, compared with the simulated AFM images used to train the network, we did not expect success. Surprisingly, however, the algorithm was able to identify chains in both types of images (Figure 5). The ability to detect these chains was able to be improved by additional image processing steps such as smoothing of the thick globular EM chains and combination of the non-background categories for the higher contrast, lower noise fluorescence images. These additions can be optionally toggled in the software. Detailed examination of the backbone contours reveals excess curvature that is introduced by the SmarTrace portion of the routine (e.g. Fig. 4a, bottom). This may arise from the particulate nature of the backbone in EM images and from the significantly different chain cross-section profile in the fluorescence images compared with AFM. Nonetheless, chains are correctly identified by the automated algorithm. We have also implemented AutoSmarTrace to determine persistence lengths of simulated molecular motor trajectories (27), demonstrating an additional application of this algorithm.

**Figure 5.**
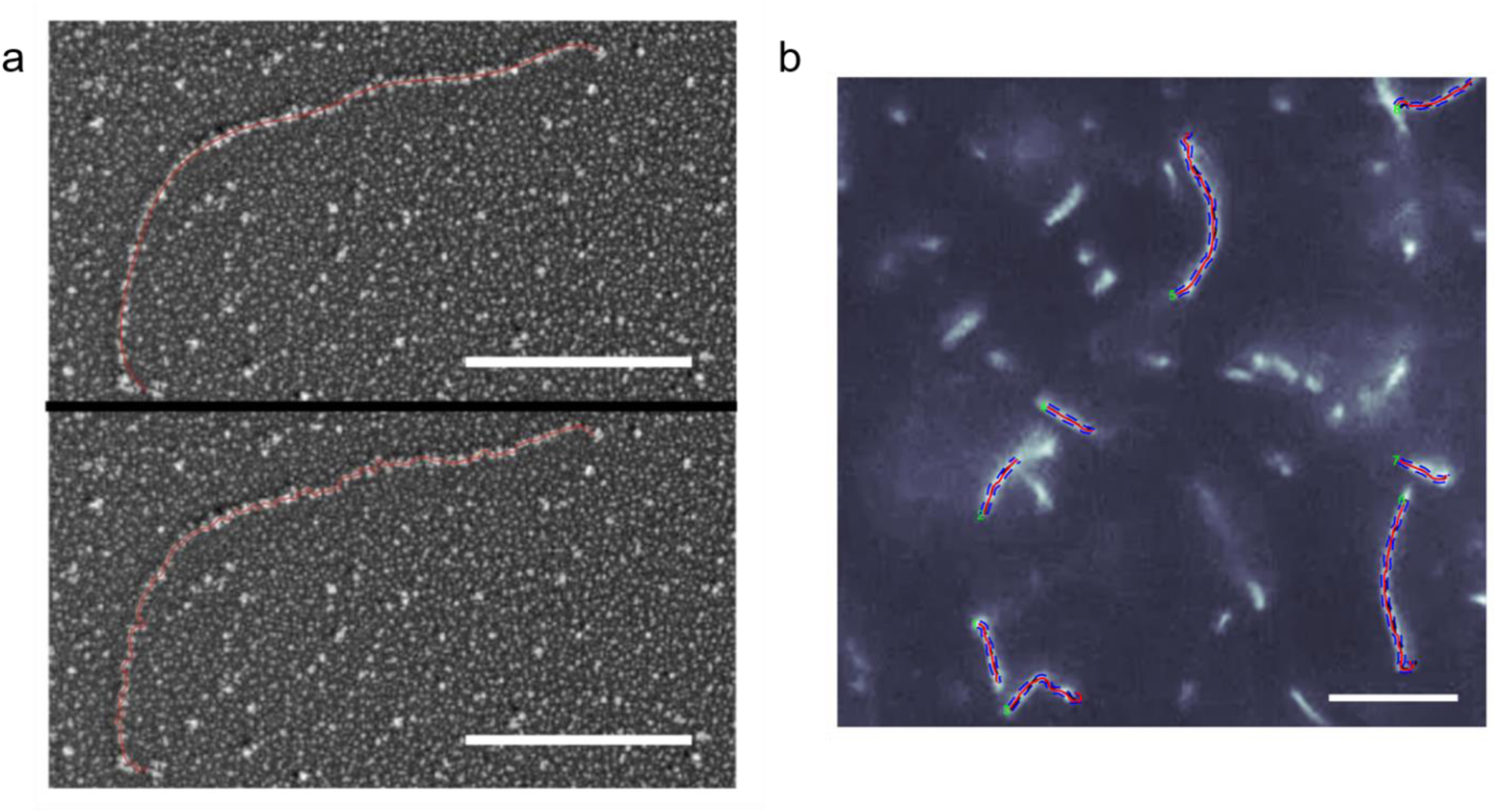
AutoSmarTrace can identify filament contours in images from different sources. (a) Electron microscopy image of type III pN-collagen. Top: red line is the initial contour spline output by the neural net + image processing portion of the algorithm, and fed as input to SmarTrace. Bottom: red line is the backbone contour identified by AutoSmarTrace. Scale bar is 100 nm. (b) Fluorescence microscopy image of actin filaments in solution. Magenta lines are the backbone contours identified by AutoSmarTrace, with blue dashed lines indicating chain widths. Scale bar is 10 μm. Image extracted from Supporting Video S1 of ref. (29).

## Discussion

Tracing filaments in AFM images can be a time-consuming process and may introduce user bias in chain selection. Here, we have shown that a machine-learning algorithm is able to correctly identify coordinates of filaments in simulated and experimental AFM images. By comparing the values of persistence length obtained by AutoSmarTrace with the ground truth used to produce simulated chain images, we validated the success of this algorithm for chains with persistence lengths ranging from 30-200 nm. As stated above, for chains with persistence lengths below ∼110 nm, analysis of tangent vector correlation is better able to recover the ground truth, while from 110-200 nm, both tangent vector correlation and mean-squared distance analysis accurately recover the persistence lengths (Fig. 3). Based on the excellent agreement between input and extracted persistence lengths at the higher end of the studied range, we anticipate that AutoSmarTrace should work well on stiffer filaments with persistence lengths > 200 nm.

Factors external to the automated chain identification algorithm may influence the estimates of persistence length. For example, our images had a pixel size of 4 nm (the conventional size of AFM scans). This makes short persistence lengths difficult to estimate accurately. For example, a chain with 20-nm persistence length will have significant directional changes over 4-5 pixels. That AutoSmarTrace can reliably recover persistence lengths as short as 30 nm (7.5 pixels) is quite impressive.

In the image processing portion of the algorithm, pixels identified as points of chain crossings by the neural net are excluded from chain tracing. This approach was chosen to eliminate the challenges of identifying correct chain directionality when emerging from a branch point, and has been used in prior manual chain tracing (5). Portions of chains are used for chain identification only if they extend, without overlapping with other chains, for at least 30 pixels. For more flexible, more highly curved and/or more dense depositions of filaments on surfaces, this selection process may require an increased number of images to attain sufficient chain contour segments for flexibility analysis.

To assess performance when tracing experimental images, we compared estimates of persistence lengths of collagen proteins via manual and automated implementations of SmarTrace (6, 7), and also benchmarked by comparing to estimates of DNA’s persistence length on images recorded and analysed by another group (5). These comparisons show the ability of AutoSmarTrace to provide consistent persistence lengths (Fig. 4). The only sample in which automatic and manual tracing led to significantly different estimates of persistence length was type IV collagen. This discrepancy likely arises from the heterogeneous structure of this network-forming collagen, compared with the fibril-forming types I and III collagens. It has been shown that type IV has an inhomogeneous flexibility profile: in particular, the persistence length is significantly lower (∼10 nm) towards the N-terminus of the protein (7, 28). Because of this, and avoidance of chain overlap in chain selection, it may be that more rigid portions of collagen IV chains are retained by the chain selection algorithm, while more flexible regions have a greater likelihood of exclusion from further analysis. Thus, filaments containing regions of high flexibility (persistence lengths on the order of a few pixels in size) may present challenges to AutoSmarTrace with the current chain selection criteria in place. For filaments with homogeneous flexibility, or whose flexibility is above 30 nm throughout the chain, there should be no selection bias towards incomplete segments of chain and AutoSmarTrace should be able to extract the correct net chain flexibility.

Two key benefits of automated chain tracing are savings of user time, and consistency of trace results. Manual tracing can lead to varying persistence length results based on the user selected points, chain selection, and subjective interpretation of a good trace. AutoSmarTrace will output identical traces each time for a given image. There may be a chain selection bias inherent to the program, but this bias will be the same across multiple runs and different users. The ability to automate this chain-tracing workflow should prove particularly beneficial for large-scale imaging applications, such as video-based analysis of dynamical filament flexibility.

The outputs of the automated chain identification software here are fed into the refined chain tracing algorithm we previously devised, dubbed SmarTrace. However, chain coordinates could alternatively be used as a starting guess (Fig. 4a, top) and provided to other tracing algorithms in place of user-required chain selection. We expect the chain-identification capabilities of AutoSmarTrace to provide greater throughput and reproducibility to analysis of filament conformations imaged using techniques that introduce different cross-sectional profiles than AFM, such as electron microscopy or fluorescence microscopy (Fig. 4).

We have shown that machine learning can provide an excellent approach to determination of biopolymer flexibility from AFM images. AutoSmarTrace identifies valid regions of chains in images, and feeds their coordinates to a refined tracing software, which subsequently determines persistence length (and curvature, where appropriate) of filaments. This approach, and others that utilize the power of computers to be trained for automated image analysis, should prove highly beneficial for increasing throughput and reliability of biopolymer characterization founded on single-molecule imaging approaches.

## Acknowledgements

Funding for this project was provided by a Discovery Grant from the Natural Sciences and Engineering Research Council of Canada (NSERC). We acknowledge many valuable discussions on image analysis with Naghmeh Rezaei, who devised and wrote the original SmarTrace algorithm. We are grateful to Daniel Sloseris for beta-testing AutoSmarTrace and testing its applicability to fluorescence image characterization. We thank Yoshihiro Ishikawa (University of California, San Francisco) for the samples of bovine pN-III collagen; Douglas R. Keene (Micro-Imaging Center, Shriners Hospitals for Children, Portland, Oregon) for recording and sharing the electron microscopy images of bovine pN-III collagen; and Patrick Heenan and Tom Perkins (JILA and University of Colorado Boulder) for sharing the AFM images of DNA.

## Supporting Information

Supporting Video shows a real-time screen capture of AutoSmarTrace chain identification and tracing in an AFM image of type I collagen chains.

## Notes

https://github.com/FordeLab/AutoSmarTrace

### Competing Interest Statement

The authors have declared no competing interest.

### Summary of Updates

Added supporting video showing AutoSmarTrace in action.

## References

1. Kirkness, M. W. H., K. Lehmann, and N. R. Forde. 2019. Mechanics and structural stability of the collagen triple helix. Current Opinion in Chemical Biology 53:98–105.

2. Chang, S.-W., Sandra J. Shefelbine, and Markus J. Buehler. 2012. Structural and Mechanical Differences between Collagen Homo- and Heterotrimers: Relevance for the Molecular Origin of Brittle Bone Disease. Biophysical Journal 102(3):640–648.

3. Gautieri, A., F. S. Passini, U. Silván, M. Guizar-Sicairos, G. Carimati, P. Volpi, M. Moretti, H. Schoenhuber, A. Redaelli, M. Berli, and J. G. Snedeker. 2017. Advanced glycation end-products: Mechanics of aged collagen from molecule to tissue. Matrix Biology 59:95–108.

4. Rivetti, C., M. Guthold, and C. Bustamante. 1996. Scanning Force Microscopy of DNA Deposited onto Mica: Equilibration versus Kinetic Trapping Studied by Statistical Polymer Chain Analysis. Journal of Molecular Biology 264(5):919–932.

5. Heenan, P. R., and T. T. Perkins. 2019. Imaging DNA Equilibrated onto Mica in Liquid Using Biochemically Relevant Deposition Conditions. ACS Nano 13(4):4220–4229.

6. Rezaei, N., A. Lyons, and N. R. Forde. 2018. Environmentally Controlled Curvature of Single Collagen Proteins. Biophysical Journal 115(8):1457–1469.

7. Al-Shaer, A., A. Lyons, Y. Ishikawa, B. G. Hudson, S. P. Boudko, and N. R. Forde. 2020. Sequence-dependent mechanics of collagen reflect its structural and functional organization. bioRxiv:2020.2009.2027.315929.

8. Brangwynne, C. P., G. H. Koenderink, E. Barry, Z. Dogic, F. C. MacKintosh, and D. A. Weitz. 2007. Bending Dynamics of Fluctuating Biopolymers Probed by Automated High-Resolution Filament Tracking. Biophysical Journal 93(1):346–359.

9. Valdman, D., Paul J. Atzberger, D. Yu, S. Kuei, and Megan T. Valentine. 2012. Spectral Analysis Methods for the Robust Measurement of the Flexural Rigidity of Biopolymers. Biophysical journal 102(5):1144–1153.

10. Faas, F. G. A., B. Rieger, L. J. van Vliet, and D. I. Cherny. 2009. DNA Deformations near Charged Surfaces: Electron and Atomic Force Microscopy Views. Biophysical Journal 97(4):1148–1157.

11. Lamour, G., J. Kirkegaard, H. Li, T. Knowles, and J. Gsponer. 2014. Easyworm: an open-source software tool to determine the mechanical properties of worm-like chains. Source Code Biol. Med. 9(1):16.

12. Usov, I., and R. Mezzenga. 2015. FiberApp: An Open-Source Software for Tracking and Analyzing Polymers, Filaments, Biomacromolecules, and Fibrous Objects. Macromolecules 48(5):1269–1280.

13. Beton, J. G., R. Moorehead, L. Helfmann, R. Gray, B. W. Hoogenboom, A. P. Joseph, M. Topf, and A. L. B. Pyne. 2020. TopoStats - an automated tracing program for AFM images. bioRxiv:2020.2009.2023.309609.

14. Rezaei, N. 2016. Mechanical Studies of Single Collagen Molecules Using Imaging and Force Spectroscopy. In Department of Physics. Simon Fraser University, Burnaby, BC, Canada.

15. Nichols, J. A., H. W. Herbert Chan, and M. A. B. Baker. 2019. Machine learning: applications of artificial intelligence to imaging and diagnosis. Biophysical Reviews 11(1):111–118.

16. Moen, E., D. Bannon, T. Kudo, W. Graf, M. Covert, and D. Van Valen. 2019. Deep learning for cellular image analysis. Nature Methods 16(12):1233–1246.

17. Mockl, L., A. R. Roy, and W. E. Moerner. 2020. Deep learning in single-molecule microscopy: fundamentals, caveats, and recent developments Invited. Biomed. Opt. Express 11(3):1633-1661. Review.

18. Zhang, P., S. Liu, A. Chaurasia, D. Ma, M. J. Mlodzianoski, E. Culurciello, and F. Huang. 2018. Analyzing complex single-molecule emission patterns with deep learning. Nature Methods 15(11):913–916.

19. Möckl, L., A. R. Roy, P. N. Petrov, and W. E. Moerner. 2020. Accurate and rapid background estimation in single-molecule localization microscopy using the deep neural network BGnet. Proceedings of the National Academy of Sciences 117(1):60–67.

20. Xu, J., G. Qin, F. Luo, L. Wang, R. Zhao, N. Li, J. Yuan, and X. Fang. 2019. Automated Stoichiometry Analysis of Single-Molecule Fluorescence Imaging Traces via Deep Learning. Journal of the American Chemical Society 141(17):6976–6985.

21. MATLAB and Statistical Toolbox Release 2018b (The MathWorks, Inc., Natick, Massachusetts, United States).

22. Li, X., W. Lehman, and S. Fischer. 2010. The relationship between curvature, flexibility and persistence length in the tropomyosin coiled-coil. Journal of Structural Biology 170(2):313–318.

23. Gentle, J. E. 2003. Random Number Generation and Monte Carlo Methods. Springer-Verlag, New York.

24. Marin-Gonzalez, A., C. L. Pastrana, R. Bocanegra, A. Martín-González, J. G. Vilhena, R. Pérez, B. Ibarra, C. Aicart-Ramos, and F. Moreno-Herrero. 2020. Understanding the paradoxical mechanical response of in-phase A-tracts at different force regimes. Nucleic Acids Research 48(9):5024–5036.

25. Timpl, R., R. W. Glanville, H. Nowack, H. Wiedemann, P. P. Fietzek, and K. Kühn. 1975. Isolation, Chemical and Electron Microscopical Characterization of Neutral-Salt-Soluble Type III Collagen and Procollagen from Fetal Bovine Skin. Biological Chemistry 356(2):1783–1792.

26. Sakai, L. Y., and D. R. Keene. 1994. Fibrillin: Monomers and microfibrils. Methods in Enzymology. Academic Press, pp. 29–52.

27. Korosec, C. S., L. Jindal, M. Schneider, I. Calderon de la Barca, M. J. Zuckermann, N. R. Forde, and E. Emberly. 2021. Substrate stiffness tunes the dynamics of polyvalent rolling motors. Soft Matter. 10.1039/D0SM01811B.

28. Hofmann, H., T. Voss, K. Kuhn, and J. Engel. 1984. Localization Of Flexible Sites In Thread-Like Molecules From Electron-Micrographs - Comparison Of Interstitial, Basement-Membrane And Intima Collagens. Journal Of Molecular Biology 172(3):325–343.

29. Takatsuki, H., E. Bengtsson, and A. MÅnsson. 2014. Persistence length of fascin-cross-linked actin filament bundles in solution and the in vitro motility assay. Biochimica et Biophysica Acta (BBA) - General Subjects 1840(6):1933–1942.

