## Supplementary figures and images for "AutoSmarTrace: Automated Chain Tracing and Flexibility Analysis of Biological Filaments"

### Supporting Video

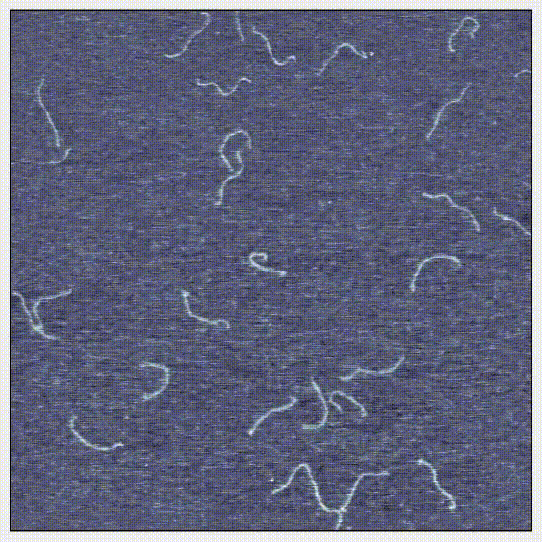
